# Machine learning algorithms can predict tail biting outbreaks in pigs using feeding behaviour records

**DOI:** 10.1101/2021.05.11.443554

**Authors:** Catherine Ollagnier, Claudia Kasper, Anna Wallenbeck, Linda Keeling, Giuseppe Bee, Siavash A. Bigdeli

**Affiliations:** Swine Research Unit, Agroscope, Posieux, Fribourg, Switzerland; Animal GenoPhenomics, Agroscope, Posieux, Fribourg, Switzerland; Department of Animal Environment and Health, Swedish University of Agricultural Sciences, Ultuna, Uppsala, Sweden; Edge Vision and AI, CSEM, Neuchâtel, Neuchâtel, Switzerland

## Abstract

Tail biting is a damaging behaviour that impacts the welfare and health of pigs. Early detection of precursor sig outbreaks, using feeding behaviour data recorded by an electronic feeder. Prediction capacities of seven machine learning algorithms (Generalized Linear Model with Stepwise Feature Selection, random forest, Support Vector Machines with Radial Basis Function Kernel, Bayesian Generalized Linear Model, Neural network, K-nearest neighbour, and Partial Least Squares Discriminant Analysis) were evaluated from daily feeding data collected from 65 pens originating from two herds of grower-finisher pigs (25-100kg), in which 27 tail biting events occurred. Data were divided into training and testing data in two different ways, either by randomly splitting data into 75% (training set) and 25% (testing set), or by randomly selecting pens to constitute the testing set. In the first data splitting, the model is regularly updated with previous data from the pen, whereas in the second data splitting, the model tries to predict for a pen that it has never seen before. The K-nearest neighbour algorithm was able to predict 78% of the upcoming events with an accuracy of 96%, when predicting events in pens for which it had previous data. The detection of events for unknown pens was less sensitive, and the neural network model was able to detect 14% of the upcoming events with an accuracy of 63%. Our results indicate that machine learning models can be considered for implementation into automatic feeder systems for real-time prediction of tail biting events.

## Introduction

Tail biting (TB) is abnormal behaviour in pigs that is thought to have a multi-factorial origin. A lack of enrichment material, unfavourable environmental conditions, an unbalanced diet, or a poor health status could trigger it. In addition to the welfare and ethical concerns associated with this cannibalistic behaviour, TB events cause pain, trigger infections, impair growth and devalue the carcasses [1-5].

Routine tail docking is prohibited in Switzerland [6] and in the EU [7], and farmers are asked to set up measures to prevent TB outbreaks. One strategy is to pinpoint the farm-specific risk factors for TB and to find solutions to reduce them [4, 6]. Another strategy is to monitor animals’ behaviours to detect early signs of forthcoming outbreaks [8-10]. Early identification of TB indicators is important for efficient intervention. The behavioural monitoring can be done at the pen and at the individual animal level. Identification at the individual animal level can support preventive measures such as removing the biter or the bitten pigs. Observations at the pen level are more efficient to detect the TB event [3].

To date, only a few behavioural indicators were studied at the pen level. Early indicators— such as changes in activity levels, tail posture, changes in exploratory behaviour, and drinking and feeding behaviours—can be observed up to 63 days before outbreaks occur, but observations are sometimes inconsistent. For instance, Statham et al. reported that pigs spend less time lying and more time standing and sitting within four days before an outbreak [11], but Wedin et al. did not observe this difference in postures [10]. Zonderland et al. reported decreasing exploratory behaviour six days before TB events [12], whereas Statham et al. observed increasing environmental manipulations one day before [11]. In contrast, Ursinus et al. did not observe any change in explorative behaviour before TB events [13]. Larsen et al. detected a change in activity and object manipulation within the 7 days before an event [14]. A lower tail position seems also to indicate an outbreak, and several authors have reported an increased incidence of tucked or hanging tails in pens before and during TB outbreaks [8-11, 15]. Using automated analysis of camera recordings, D’Eath et al. and Liu et al. detected low tail posture, which was positively associated with more tail damage [8, 16]. Nonetheless, the tail posture may not specifically indicate a TB event, since low tail posture has also been associated with negative emotional responses in pigs [10], which could be caused by other factors like sickness. All of the above studies described behavioural changes when comparing a control (CTL) pen to a TB pen, which is the first step in developing of early detection of TB event. The statistical analyses identify significant changes in behaviour before and during a TB outbreak, but none of the authors attempted to use the detected differences to predict upcoming events. In addition, the behavioural traits monitored in the previous studies require regular observations or additional material (camera) to detect changes in behaviour, which is either time consuming or costly.

Nowadays, more and more pig farms are equipped with electronic feeding systems. The technology offers individually tailored feeding, reduces pigs feed usage, improves health and welfare, and reduces farm workload. Automatic pig feeding systems bring increased efficiency, convenience and control to the feeding process. Electronic feeding systems with single-spaced feeders also enable automatic monitoring of the feeding behaviours of each individual. Recording the identity of the pig, feeder entry and exit times and the amount of food consumed allows the calculation of the frequency of feeder visits per day, feeding rates, mean feeder occupation time, mean food intake per feeder visit, total food intake and total feeder duration per day for each pig. In 1994, Young and Lawrence found that pigs housed in groups and fed from automatic feeders showed a temporal pattern of feeding behaviour [17]. They also suggested that the feeding behaviour might be altered by social conditions. It has been later described that changes in feeding behaviours with automatic feeders were associated with negative events like aggressive behaviour or disrupted social dynamics [8]. If a TB event can be predicted from behaviour, as postulated by Statham et al. [11], then data from electronic feeders could be used to monitor in real time the feeding behaviour of the pigs. In fact, the feeding behaviour of pigs assessed by electronic feeders appears to change before a TB event. Some studies describe changes in feeding behaviours before TB events [18-20]. These findings suggest that feeding behaviours recorded by electronic feeders could be a valid tool to detect early signs of a TB event. Indeed, Maselyne et al. developed an online warning system for individual fattening pigs based on their feeding pattern [21]. This study investigated whether abnormal changes in the feeding pattern can be detected automatically and used as an (early) indicator for health, welfare and productivity problems of an individual animal. They observed the number of feeder visits per day and the average time interval between two visits and determined a threshold above which the behaviour was considered abnormal. Every pig was categorised each day as ‘green’ (globally healthy), ‘orange’ or ‘red’ status (the latter including severe infection of the tail). However, the authors worked at the individual pig level and did not focus on TB detection at the pen level.

Different authors attempted to predict three behavioural changes (pen fouling, diarrhoea and TB) using multiple data types extracted from the pen [22-24]. A multivariate dynamic model and/or machine neural network and Bayesian ensemble were created by combining information from the drinking and feeding behaviours of pigs and the pen’s environmental conditions. In these articles, the authors acknowledged that feed and water consumption are highly correlated [22] and that changes in water consumption are better predictors of behavioural changes than environmental parameters [24]. Due to a lack of TB data during the period of Jensen et al.’s analysis, the researchers were unable to predict TB event [23]. The aforementioned studies were limited by the fact that they rely on water/climate sensors, which are not routinely installed in farms. Further, the authors did not address whether their model could be generalized to another farm data set.

In our study, we used feeding behaviour data paired with machine learning (ML) algorithms to predict TB outbreaks in real time. The study’s objectives are: 1) assessment of the impact of the data framework on TB detection; 2) implementation and evaluation of the proposed framework on two different farm datasets; 3) assessment of a data-independent model; 4) evaluation of the framework’s impact on TB detection.

In summary, the contributions of our research are:

1. Provide a new data framework to allow a ML approach to predict TB using feeding behaviour data;
2. Demonstrate that Machine Learning Models —Generalized Linear Model with Stepwise Feature Selection (glmnet), random forest (rf), Support Vector Machines with Radial Basis Function Kernel (svmRadial), Bayesian Generalized Linear Model (bayesglm), Neural network (nn), K-nearest neighbour (kNN), and Partial Least Squares Discriminant Analysis (pls)— can predict TB events using pigs’ feeding behaviours at the pen level with the new data framework;
3. Simulate two conditions: one where the model has access to previous data of the pen, and another where the model makes predictions in one pen, based on data from other pens;
4. Achieve a prediction of 70-80% of the upcoming TB events with a specificity of >99% (rf and kNN models), when the model has access to previous data of the pen, and
5. Evaluate and compare prediction performances in two different farm conditions.

A TB monitoring tool would open up new opportunities for the farmer to take targeted action in specific pens to prevent the TB event. Being able to prevent TB would serve the welfare of the animals and provide economic benefits to the farmer. Since the tool requires only data that are already available from pig farms equipped with automatic feeders, it could be easily implemented in commercial practice as an additional management tool.

## Material and methods

### Data collection

This study analyses the feeding behaviours of two herds of grower-finisher pigs weighing between 25 and 100 kilograms. One data set originates from a testing boar station in Sweden and contains data collected from October 2004 to July 2007. The data set comes from a previous retrospective study that Wallenbeck and Keeling published [20]. The second data set contains data from the experimental pig farm of Agroscope and comprises recordings from November 2018 to April 2020. As tail docking is prohibited in Sweden and in Switzerland, the data are from pigs with intact tails.

The Swedish data set includes data from 42 pens (21 TB and 21 CTL) of boars (purebred Yorkshire, Landrace or Hampshire) recorded 70 days before and after the TB date. Boars were housed in groups of 7 to 14 animals per pen. Each pen measured 15.7 m and had a slatted floor and plain resting area. All pigs had *ad libitum* access to the pelleted feed, which was optimised according to the Swedish nutrition norms for fattening pigs [25]. Water was provided *ad libitum* and straw was offered daily.

The Swiss data set consisted of 23 pens (six TB and 17 CTL) of females and castrated male pigs (Swiss Large White), recorded 100 days before and after the TB date. Twenty pens (18 m) contained 11 to 15 pigs each and were equipped with two automatic feeders; three pens (78 m) were equipped with eight automatic feeders for 31 to 55 pigs each. All pens had straw in racks and sawdust on the floor. Water was available *ad libitum* through nipple drinkers. The pelleted finisher diet was formulated to have 20% lower dietary crude protein and essential amino acids compared to a standard diet formulated according to the Swiss feeding recommendations for pigs [26].

For both study sites, data were collected by individual automatic feeders (ACEMO 48, Acemo, France; or MLP, Agrotronic Schauer, Austria) that recorded the number of visits to the feeder and the amount of feed consumed. The feeders were 0.6 m wide and 1.5-2.2 m long. Only one pig could enter the feeder at a time, and other pigs could not dislodge the pig feeding inside the feeder. Each pig had access to only one feeder.

A pen was assigned to the TB category if at least one pig had to be treated for tail damages. The TB date (day 0) was defined as the date at which the first treatment was recorded. For the Swedish data set, each pen in the TB category was paired to a pen in the CTL category. For the Swiss data set, all the 23 farrowing batches reared under the same housing and feeding conditions were considered for analysis.

### Definitions of the observations, analysis in time series, and missing value imputation

The frequency of daily feeder visits (DFV), the daily feed consumption (DFC), and the standard deviation of the feed consumption at each visit (StdFC) were calculated per day and per pig (Table 1). These parameters were considered as ‘observations’ to predict TB events at the pen level and were derived from the data collected by the automatic feeder.

**Table 1.** Observations used for tail biting predictions.

| Observations | Definition | Units | Abbreviated |
| --- | --- | --- | --- |
| **Frequency of daily feeder visits** | Number of visits to the feeder (from 0:00 to 23:59:59 that date) | n | DFV |
| **Daily feed consumption** | Total feed consumption (from 0:00 to 23:59:59 that date) | g | DFC |
| **Standard deviation of the feed consumption** | Daily standard deviation of the feed consumption at each visit | g | StdFC |

The time dependency of the observations was taken into account by analysing the data by groups of consecutive data points, called the ‘analysis window’. The prediction model considered the analysis window to achieve a prediction at the pen level. The analysis window was first defined to contain observations from 14 consecutive days (Fig 1). The A_date was defined as the first day of the analysis window. The analysis window slides along the timeline and the A_date defines the class of the analysis window, i.e., “TB” or “CTL”. Analysis windows from CTL pens were always classified as “CTL”, whereas analysis windows from TB pens were considered as “TB” class only between day -35 and day 10 (TB window). The analysis window of a TB pen was classified as “TB” if the A_date was inside the TB window and “CTL”, if outside the TB window. Missing values were computed using median imputation (by data set) and a principal component analysis was performed before ML analysis.

**Fig 1.**
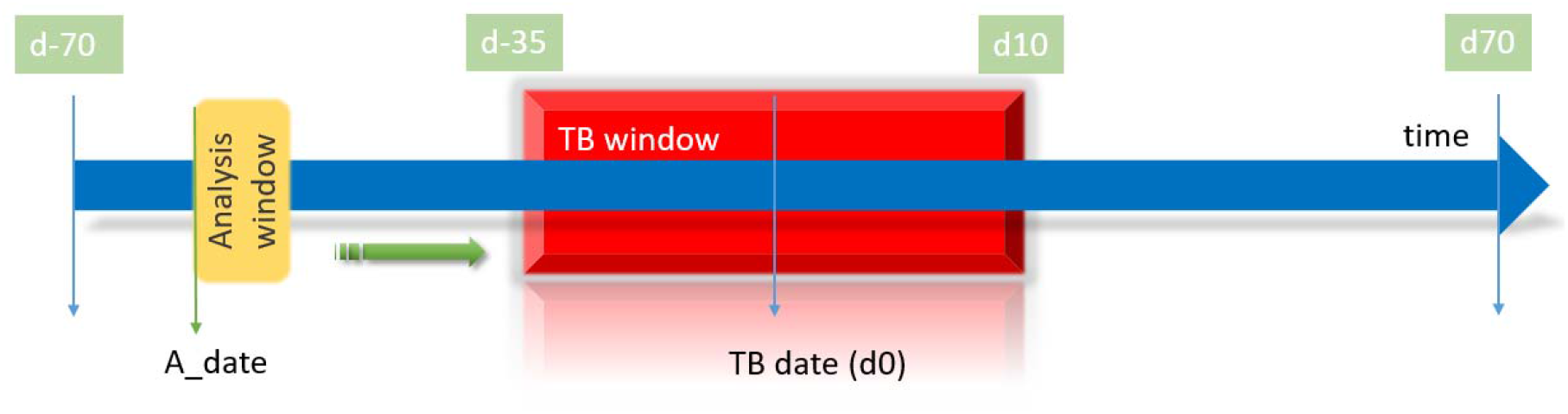
Analysis of the time dependency of the data thanks to the analysis window approach. Analysis window slides along the timeline (blue arrow). The analysis window is classified as TB class, when the A_date (first day of the analysis window) enters the TB window (orange block) and as CTL class when it is outside the TB window. Control pens are always classified as “CTL”.

In each pen, the analysis window contained observations from 10 pigs for 14 days, to standardize the size of the analysis window. Observations from 10 pigs were considered to give enough information on the pen, without creating too many missing data points, for the few occurrences that contained fewer than 10 pigs.

At the end of the data framing, each analysis window contained 420 observations [3 variables (DFV, DFC, StdFC) × 10 pigs × 14 days], and one outcome (the class of the window: TB or CTL). The Swiss and the Swedish data set were merged into a third data set, called Swedish+Swiss, to incorporate more diverse observations and further increase the model’s generalizability for unseen data (pen/country). In total, the combined data set (Swedish and Swiss data) contained 6605 analysis windows, with 5479 and 1126 CTL and TB windows, respectively. The characteristics of the TB windows compared to CTL are presented in Table 2.

**Table 2.** Characteristics of the data sets.

|  |  | Swedish |  | Swiss |  |
| --- | --- | --- | --- | --- | --- |
| Observations | Statistics | CTL | TB | CTL | TB |
| DFC<sup>1</sup> | Mean | 2337.1* | 2005.7 | 2282.3* | 2438.8 |
|  | SD | 757.4 | 708.6 | 600.2 | 678.2 |
| DFV<sup>2</sup> | Mean | 24.9* | 24.0 | 12.3* | 12.0 |
|  | SD | 19.4 | 18.0 | 6.0 | 7.2 |
| StdFC<sup>3</sup> | Mean | 128.6* | 113.8 | 166.8* | 195.6 |
|  | SD | 85.9 | 71.2 | 96.3 | 129.7 |

\*Significant difference between CTL and TB analysis window classes (p<0.0001)
<sup>1</sup> DFC: frequency of daily feeder visits
<sup>2</sup> DFV: daily feed consumption
<sup>3</sup> StdFC: standard deviation of the feed consumption at each visit.

### Models

All data were analysed with R3.6.3, using the caret package to build ML models [27]. Commonly used classification ML methods were first tested on all three data sets (i.e., Swedish, Swiss and Swedish+Swiss) [28, 29]. Table 3 presents a list of common ML methods considered and implemented with R packages in this study with binary outcome.

**Table 3.** The seven machine learning methods used to predict tail biting events from feeding behaviour data.

| Machine Learning | R Package | Function |
| --- | --- | --- |
| Generalized Linear Model with Stepwise Feature Selection | glmnet | glmnet |
| Random forest | ranger | rf |
| Support Vector Machines with Radial Basis Function Kernel | kernlab | svmRadial |
| Bayesian Generalized Linear Model | arm | bayesglm |
| Neural network | nnet | nn |
| K-nearest neighbour | caret | kNN |
| Partial Least Squares Discriminant Analysis | pls | pls |

For each data set, predictive models were first trained on a subset of data (training set), and the models’ performances on this training set were then compared. The predictive performances of the models were further compared using the unseen data (test set). The test set contained either new unseen analysis windows (cross-validation (CV) approach) or new pens (leave one out cross-validation (LOOP) approach) [30]. The LOOP approach gives estimate metrics that are valuable when a new pen is presented to the model for prediction, as there is no pen overlapping between the training and testing data sets. This represents the situation where the farmer tries to predict a TB event in a hitherto unknown pen. The CV resampling approach predicts TB events based on data previously recorded in the pen. This approach is correct when the model can be continuously updated with previous records of the pen so that the prediction model already knows the feeding behaviour of the pen and tries to classify the analysis window of the testing set based on previous knowledge of this pen.

### Model evaluation: performances metrics

This is a classification problem with binary outcomes (TB or CTL), and performances of the models should be assessed on parameters calculated with a confusion matrix [31]. Performance metrics definitions and confusion matrix are presented in Table 4. The sensitivity (rate of predicted TB class given the actual TB class) assesses the capacity of the model to detect an upcoming TB event. The positive predictive value (PPV) evaluates the capacity of the model to correctly predict a TB class. All models were optimized to maximize the sensitivity, as this study aimed to detect early warnings of TB events. The specificity (rate of predicted CTL given the actual CTL class) assesses the capacity of the model to detect a normal behaviour. The kappa statistic assesses how the model outperforms a random model that simply always predicts “CTL”. According to Landis and Koch, a kappa of 0-0.20 is slight, 0.21-0.40 is fair, 0.41-0.60 is moderate, 0.61-0.80 is substantial, and 0.81-1 is almost perfect [32]. The *p*-value assesses the statistical significance of the difference in accuracy between the random model and the tested model.

**Table 4.** The confusion matrix and performances metrics used to assess the performances of the models. A confusion matrix was applied to evaluate the prediction performances of the ML models for this classification problem with a binary outcome (tail biting, “TB” or control “CTL” class). The definitions of the performances metrics are presented.

|  |  | Actual |  |
| --- | --- | --- | --- |
|  |  | TB | CTL |
| Predicted | TB | **TP**<br>**True positive**<br>predicted TB class that are actually TB class | **FP**<br>**False-positive**<br>predicted TB class that are actually CTL class |
|  | CTL | **FN**<br>**False-negative**<br>TB class not detected by the model (predicted as CTL class) | **TN**<br>**True negative**<br>predicted CTL class that are actually CTL class |
| **Performances Metrics**<br>(TP, FP FN and TN are defined above) |  |  |  |
| | | $Sensitivity = \frac{TP}{TP + FN}$ | (1) |
| | | $Positive Predicted Value (PPV) = \frac{TP}{TP + FP}$ | (2) |
| | | $Specificity = \frac{TN}{TN + FP}$ | (3) |
| | | $Accuracy = \frac{TN + TP}{TN + FP + TP + FN}$ | (4) |
|  |  | *P-value*: statistical significance of the difference with a random model always predicting CTL class. |  |
| | | $K = \frac{p_o - p_e}{1 - p_e}$ where $p_o$ is the observed accuracy, and $p_e$ is the expected accuracy of a random model always predicting CTL class. | (5) |

## Results

### Model performances

Table 5 presents the model performances on the three training data sets. All models performed significantly better than the random prediction model (that simply always predict CTL), with kappas ranging from 0.30 to 1.00. Even if the criteria for optimization was the sensitivity, this performance criterion was always lower or equal to the specificity, which is most likely due to the imbalance between the numerous CTL and the rare TB classes.

**Table 5.**
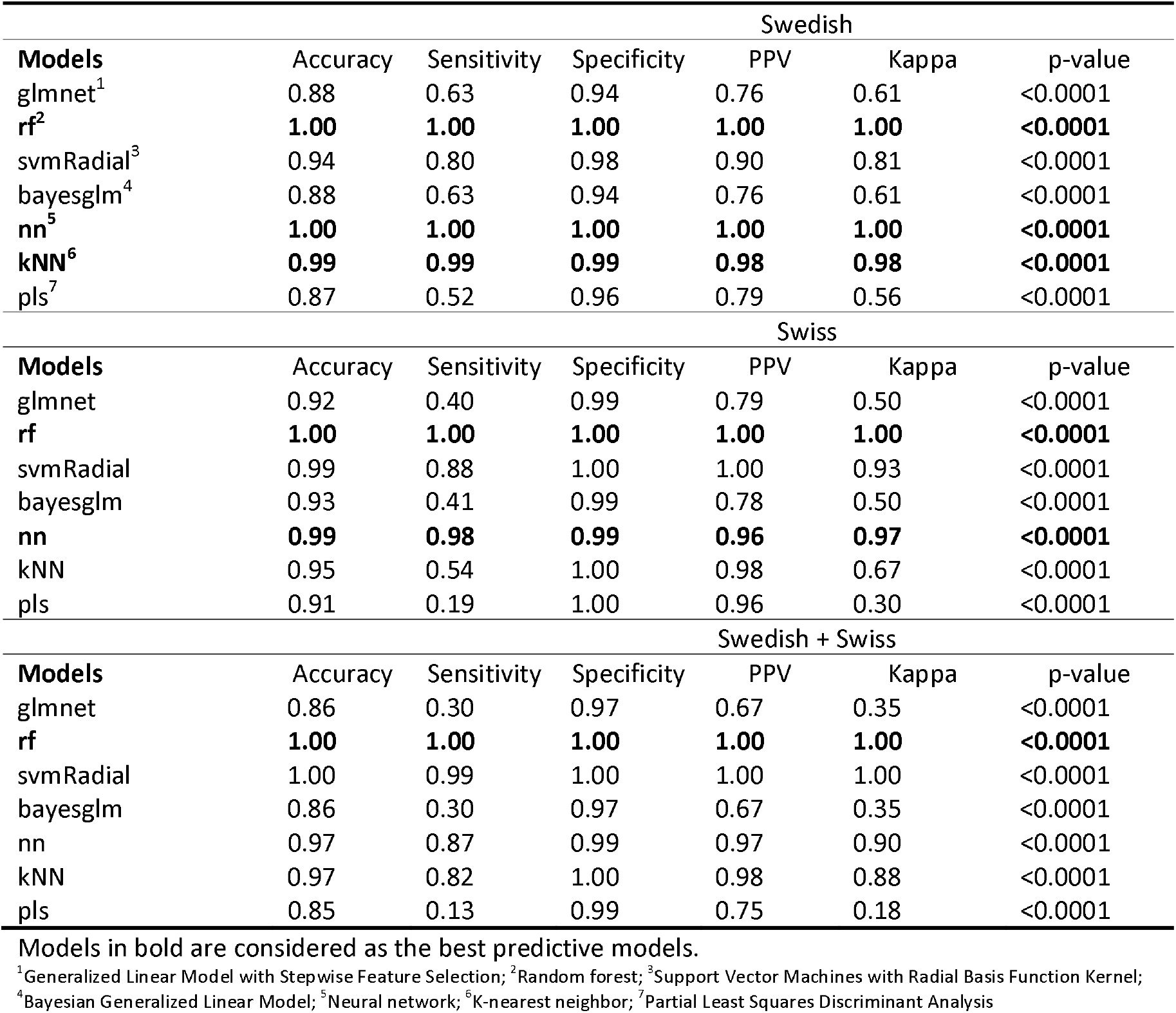
Models performances on <u>training data sets</u> for the Swiss, Swedish and Swedish+Swiss data sets.

| Swedish |  |  |  |  |  |  |
| --- | --- | --- | --- | --- | --- | --- |
| Models | Accuracy | Sensitivity | Specificity | PPV | Kappa | p-value |
| glmnet<sup>1</sup> | 0.88 | 0.63 | 0.94 | 0.76 | 0.61 | <0.0001 |
| **rf<sup>2</sup>** | **1.00** | **1.00** | **1.00** | **1.00** | **1.00** | **<0.0001** |
| svmRadial<sup>3</sup> | 0.94 | 0.80 | 0.98 | 0.90 | 0.81 | <0.0001 |
| bayesglm<sup>4</sup> | 0.88 | 0.63 | 0.94 | 0.76 | 0.61 | <0.0001 |
| **nn<sup>5</sup>** | **1.00** | **1.00** | **1.00** | **1.00** | **1.00** | **<0.0001** |
| **kNN<sup>6</sup>** | **0.99** | **0.99** | **0.99** | **0.98** | **0.98** | **<0.0001** |
| pls<sup>7</sup> | 0.87 | 0.52 | 0.96 | 0.79 | 0.56 | <0.0001 |
| Swiss |  |  |  |  |  |  |
| Models | Accuracy | Sensitivity | Specificity | PPV | Kappa | p-value |
| glmnet | 0.92 | 0.40 | 0.99 | 0.79 | 0.50 | <0.0001 |
| **rf** | **1.00** | **1.00** | **1.00** | **1.00** | **1.00** | **<0.0001** |
| svmRadial | 0.99 | 0.88 | 1.00 | 1.00 | 0.93 | <0.0001 |
| bayesglm | 0.93 | 0.41 | 0.99 | 0.78 | 0.50 | <0.0001 |
| **nn** | **0.99** | **0.98** | **0.99** | **0.96** | **0.97** | **<0.0001** |
| kNN | 0.95 | 0.54 | 1.00 | 0.98 | 0.67 | <0.0001 |
| pls | 0.91 | 0.19 | 1.00 | 0.96 | 0.30 | <0.0001 |
| Swedish + Swiss |  |  |  |  |  |  |
| Models | Accuracy | Sensitivity | Specificity | PPV | Kappa | p-value |
| glmnet | 0.86 | 0.30 | 0.97 | 0.67 | 0.35 | <0.0001 |
| **rf** | **1.00** | **1.00** | **1.00** | **1.00** | **1.00** | **<0.0001** |
| svmRadial | 1.00 | 0.99 | 1.00 | 1.00 | 1.00 | <0.0001 |
| bayesglm | 0.86 | 0.30 | 0.97 | 0.67 | 0.35 | <0.0001 |
| nn | 0.97 | 0.87 | 0.99 | 0.97 | 0.90 | <0.0001 |
| kNN | 0.97 | 0.82 | 1.00 | 0.98 | 0.88 | <0.0001 |
| pls | 0.85 | 0.13 | 0.99 | 0.75 | 0.18 | <0.0001 |

Models in bold are considered as the best predictive models.
<sup>1</sup>Generalized Linear Model with Stepwise Feature Selection; <sup>2</sup>Random forest; <sup>3</sup>Support Vector Machines with Radial Basis Function Kernel;
<sup>4</sup>Bayesian Generalized Linear Model; <sup>5</sup>Neural network; <sup>6</sup>K-nearest neighbor; <sup>7</sup>Partial Least Squares Discriminant Analysis

### Model prediction performances

Tables 6 and 7 summarize the performances of the models to predict unseen analysis windows (CV) or unseen pens (LOOP), respectively. For the Swedish+Swiss data set, the performances of the model on CV and LOOP were assessed on the combined testing set (Swedish+Swiss) and on subsets of the Swedish or Swiss data set separately. The RF model showed the best predictive performances on both the unseen analysis windows and the unseen pens, with an average accuracy of 84% and a sensitivity of 38% on all data sets. Predictive performances were higher on unseen analysis windows than on unseen pens, and always lower than the model performances on the training set. In the Swedish data set, the predictive performances of the RF model on unseen pens were poor (kappa<0). Models trained on the Swedish+Swiss data set showed a poorer performance (low kappa) in predicting new analysis windows of a Swedish or Swiss subset data set than the same model trained on the Swedish or Swiss data sets individually. Lower kappas were also obtained for prediction on new pens—except for glmnet, bayesglm, and pls models—that had improved predictive performances (kappa) on the Swiss subset.

**Table 6.** Performances of models to predict <u>unseen analysis windows</u> [Cross Validation (CV) approach] of the Swedish, Swiss and Swedish+Swiss testing data sets.

| Swedish |  |  |  |  |  |  |
| --- | --- | --- | --- | --- | --- | --- |
| Models | Accuracy | Sensitivity | Specificity | PPV | Kappa | p value |
| glmnet<sup>1</sup> | 0.80 | 0.45 | 0.90 | 0.57 | 0.38 | 0.04 |
| **rf<sup>2</sup>** | **0.95** | **0.80** | **0.99** | **0.98** | **0.85** | **<0.0001** |
| svmRadial<sup>3</sup> | 0.86 | 0.54 | 0.95 | 0.75 | 0.54 | <0.0001 |
| bayesglm<sup>4</sup> | 0.81 | 0.45 | 0.90 | 0.57 | 0.39 | 0.04 |
| nn<sup>5</sup> | 0.87 | 0.70 | 0.91 | 0.70 | 0.62 | <0.0001 |
| **kNN<sup>6</sup>** | **0.97** | **0.94** | **0.98** | **0.94** | **0.92** | **<0.0001** |
| pls<sup>7</sup> | 0.81 | 0.36 | 0.93 | 0.60 | 0.35 | - |
| Swiss |  |  |  |  |  |  |
| Models | Accuracy | Sensitivity | Specificity | PPV | Kappa | p-value |
| glmnet | 0.85 | 0.22 | 0.94 | 0.30 | 0.17 | - |
| **rf** | **0.94** | **0.50** | **0.99** | **0.95** | **0.63** | **<0.0001** |
| svmRadial | 0.91 | 0.28 | 1.00 | 0.92 | 0.39 | 0.006 |
| bayesglm | 0.85 | 0.22 | 0.93 | 0.30 | 0.17 | - |
| **nn** | **0.89** | **0.47** | **0.95** | **0.54** | **0.44** | - |
| **kNN** | **0.93** | **0.41** | **1.00** | **1.00** | **0.55** | **<0.0001** |
| pls | 0.89 | 0.11 | 0.99 | 0.56 | 0.15 | - |
| Swedish + Swiss |  |  |  |  |  |  |
| Models | Accuracy | Sensitivity | Specificity | PPV | Kappa | p-value |
| Glmnet<sup>a</sup> | 0.82 | 0.15 | 0.96 | 0.43 | 0.15 | - |
| Swedish<sup>b</sup> | 0.77 | 0.21 | 0.93 | 0.46 | 0.18 | - |
| Swiss<sup>c</sup> | 0.88 | 0.02 | 0.99 | 0.20 | 0.02 | - |
| **rf<sup>a</sup>** | **0.94** | **0.70** | **0.99** | **0.96** | **0.78** | **<0.0001** |
| Swedish<sup>b</sup> | **0.95** | **0.80** | **0.99** | **0.95** | **0.83** | **<0.0001** |
| Swiss<sup>c</sup> | **0.94** | **0.48** | **1.00** | **1.00** | **0.62** | **<0.0001** |
| svmRadial<sup>a</sup> | 0.88 | 0.38 | 0.98 | 0.80 | 0.46 | <0.0001 |
| Swedish<sup>b</sup> | 0.86 | 0.48 | 0.97 | 0.80 | 0.52 | <0.0001 |
| Swiss<sup>c</sup> | 0.90 | 0.16 | 1.00 | 0.76 | 0.23 | - |
| bayesglm<sup>a</sup> | 0.82 | 0.16 | 0.96 | 0.43 | 0.16 | - |
| Swedish<sup>b</sup> | 0.77 | 0.21 | 0.93 | 0.46 | 0.19 | - |
| Swiss<sup>c</sup> | 0.82 | 0.16 | 0.96 | 0.43 | 0.15 | - |
| **nn<sup>a</sup>** | **0.88** | **0.55** | **0.94** | **0.67** | **0.53** | **<0.0001** |
| Swedish<sup>b</sup> | **0.87** | **0.63** | **0.94** | **0.75** | **0.61** | **<0.0001** |
| Swiss<sup>c</sup> | **0.88** | **0.37** | **0.95** | **0.47** | **0.35** | - |
| **kNN<sup>a</sup>** | **0.96** | **0.78** | **1.00** | **0.98** | **0.84** | **<0.0001** |
| Swedish<sup>b</sup> | **0.96** | **0.87** | **0.98** | **0.94** | **0.88** | **<0.0001** |
| Swiss<sup>c</sup> | **0.92** | **0.32** | **1.00** | **1.00** | **0.46** | **0.0006** |
| pls<sup>a</sup> | 0.83 | 0.06 | 0.99 | 0.62 | 0.09 | - |
| Swedish<sup>b</sup> | 0.79 | 0.09 | 0.98 | 0.62 | 0.11 | - |
| Swiss<sup>c</sup> | 0.89 | 0.00 | 1.00 | 0.00 | 0.00 | - |

Models in bold are considered as the best predictive models.
<sup>1</sup>Generalized Linear Model with Stepwise Feature Selection; <sup>2</sup>Random forest; <sup>3</sup>Support Vector Machines with Radial Basis Function Kernel;
<sup>4</sup>Bayesian Generalized Linear Model; <sup>5</sup>Neural network; <sup>6</sup>K-nearest neighbour; <sup>7</sup>Partial Least Squares Discriminant Analysis
<sup>a</sup>Prediction performances of the models on testing data set containing Swedish and Swiss data.
<sup>b</sup>Prediction performances of the models on the Swedish subset of the testing data set.
<sup>c</sup>Prediction performances of the models on the Swiss subset of the testing data set.

**Table 7.**
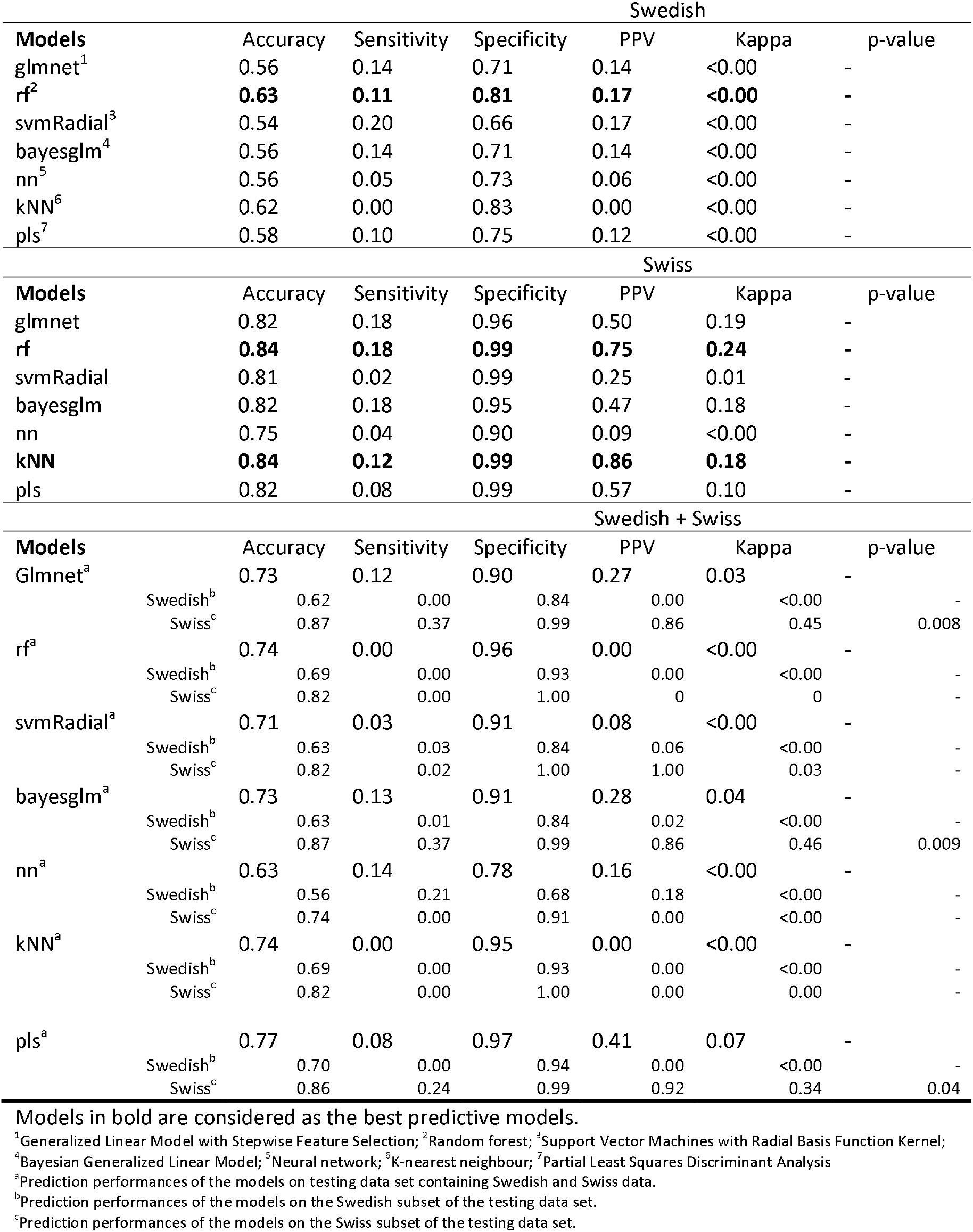
Performances of models to predict <u>unseen pens</u> [Leave one out cross-validation (LOOP) approach] of the Swedish, Swiss and Swedish+Swiss data sets.

### Impact of TB and analysis windows

We have tested the influence of the size of the analysis window and the size of the TB window on predictive performances for the Swedish data set. Accuracy of models with analysis windows of seven or 21, or with different TB windows (−49 to +10 days, or -10 to +5 days) were compared with the default windows size (analysis window: 14 days; TB window: -35 to +10 days). The RF model was chosen for this analysis, as it was the model with the best predictive performances over the three data sets and for the two (CV and LOOP) approaches.

The predictive performances of the RF model increased when the analysis window contained 21 days (instead of 14 days) or when the TB window was larger (−49 to +10 days) (Table 8). Interestingly, the RF model never predicted TB class in CTL pens, thus ML models were able to discriminate CTL pen behaviour from TB pen behaviour. In addition, the prediction of a TB event for a TB pen was almost exclusively within the TB window (high accuracy). The detection of an upcoming TB event was not better near the TB date (day 0) than at the beginning of the TB window (day -35, for default TB window size).

**Table 8.** Prediction performances of the random forest (rf) model depending on the TB and analysis window size on the Swedish data set.

| Swedish |  |  |  |  |  |  |  |  |
| --- | --- | --- | --- | --- | --- | --- | --- | --- |
| Test set | Analysis window size | TB window size | Accuracy | Sensitivity | Specificity | PPV | Kappa | p-value |
| Unseen analysis windows (CV) | 7 days | [-35;10] | 0.94 | 0.76 | 0.99 | 0.96 | 0.81 | <0.0001 |
|  |  | [-35;10] | 0.94 | 0.70 | 0.99 | 0.96 | 0.78 | <0.0001 |
|  | 14 days | [-49;10] | 0.97 | 0.88 | 0.99 | 0.97 | 0.90 | <0.0001 |
|  |  | [-10;5] | 0.95 | 0.45 | 1.00 | 0.94 | 0.58 | <0.0001 |
|  | 21 days | [-35;10] | 0.97 | 0.87 | 1.00 | 0.99 | 0.91 | <0.0001 |
| Unseen pens (LOOP) | 7 days | [-35;10] | 0.64 | 0.00 | 0.87 | 0.00 | <0.00 | - |
|  |  | [-35;10] | 0.63 | 0.11 | 0.81 | 0.17 | <0.00 | - |
|  | 14 days | [-49;10] | 0.61 | 0.19 | 0.82 | 0.35 | 0.02 | - |
|  |  | [-10;5] | 0.91 | 0.00 | 1.00 | 0.00 | 0.00 | - |
|  | 21 days | [-35;10] | 0.66 | 0.00 | 0.88 | 0.00 | <0.00 | - |

## Discussion

In this study, TB events could be detected up to 35 days in advance using an ML model that analysed feeding behaviours recorded by electronic feeders. Early indicators of TB events were more easily identified when the model had access to previous records of the pen. Thus, one should consider performing continuous analysis of the data of each pen, even in the absence of TB events. To our knowledge, this is the first time an ML algorithm is able to predict TB events in pigs using feeding behaviour data.

In the following discussion, we compare the performances of our models with those of other models that predict TB events based on other behavioural changes. In the next section, we discuss the challenge of generalizing the model to a different farm data set. Then, we conclude by presenting an interpretation of the changes in feeding behaviour associated with a TB event.

### Performances evaluation

This study obtained prediction performances comparable to studies that used drinking behaviour and climate data [22, 24], which obtained a specificity range of 44-72% and a sensitivity range of 59-100% for TB prediction. The approach taken by Larsen et al. to predict TB events deserves consideration [24]. They modelled different data sources (from drinking behaviour and climate conditions) with dynamic linear models (similar to [23]) and data were used by an artificial neural network to predict TB events, pen fouling and diarrhoea. As a parallel process, the different data sources were combined into a logistic regression model to estimate the probability of events, which was then converted to an event prediction based on a prediction threshold. Finally, these predictions were assembled in a Bayesian ensemble model to compute a final prediction. Accordingly, Domun et al. included climate ventilation system data and pig characteristics along with the same study data to compile three dynamic models and a long short-term memory neural network to forecast TB events, pen fouling, and diarrhea [22].

In the two studies cited above, data structures were different from those in the present study. Larsen et al. had a TB windows ranging from 1 to 3 days [24], while Domun et al. had an analysis window combining short-term memory (10 minutes beforehand) with long-term memory (up to 7 days) [22]. As noticed in our study, the authors observed that increasing the TB/analysis windows improved the prediction performances. The authors, however, did not include both analysisand TB windows. The analysis window provided the possibility of detecting abnormal feeding behaviour, as well as recognizing abnormal progress over 14 days. A TB window offers the opportunity to predict an impending TB outbreak as it develops, even several days before signs of tail biting damage become evident.

### Model generalisation

A system for detecting pigs’ tail biting events is difficult to develop since pigs’ behaviors tend to be complex because of the multifactorial causes of tail biting. There is no clear answer about what prediction method or pattern can be used for detecting specific events. It seems that the feeding behaviour is not only specific to a pen, but each site has its own characteristics (e.g., breeds, climates, and feed compositions). The model trained on one data set (Swedish or Swiss) was not able to detect events in the other data set, and the combination of the data sets (Swedish + Swiss) had little effect on model performances. As a result, the combined Swedish and Swiss data did not provide many advantages. It is difficult to generalize feature-based models to unseen data, as health and welfare problems often differ between herds and meaningful features are sometime hard to identify [22]. One explanation could be linked to site-specific risk factors. As acknowledged by the European Food Safety Authority in 2014, one of the difficulties in preventing TB resides in the fact that every farm is different and has its own risk factors. Prevention strategies therefore need to be designed at a farm-specific level [5]. Feeding behaviour associated with a TB event will differ depending on the chronic risk factors on the farm (breed, sex, feeding, and access to manipulable material, space, and group size). This observation was already acknowledged by Taylor et al. [33] and Valros [4], when they defined the four types of TB (two-stage, sudden-forceful, obsessive, and epidemic), associated with four putative causations. For these authors, the two-stage type is the result of chronic and moderate stress. Competition for resources is thought to cause sudden-forceful types. And generally, the epidemic type occurs after a significant change in the pig’s daily routine (e.g., food disturbance, temperature change). The obsessive type is caused by one individual pig (the tail bitter) that possibly experience long-term challenges.

### Understanding changes in feeding behaviour

Previously, Munsterhjelm et al. [16] investigated feeding behaviours of pigs 70 days before and 28 days after the TB event was detected in a pen. Feeding behaviours in TB pens were compared to matched CTL pens. Pigs in TB pens tend to visit the feeder less frequently than pigs in CTL pens. Pigs in TB pens also had less time spent at the feeder, as well as a lower daily feed consumption (DFC). This tendency persisted after the TB date, and pigs continued to spend less time in the feeder and visit less frequently, even if the difference from the control group decreased with time. Pigs also tended to eat faster (more intake per second). They concluded that the rapid change in feeding behaviour suggests that TB behaviour escalates 14 days before the TB date.

The change in feeding behaviour in the TB pen was also observed by Tessier et al. [19] during a TB outbreak in a pen. Specifically, they studied the evolution of DFV, the DFC and feeding time seven days before (pre-injury phase), seven days during (acute phase), and seven days after (recovery phase) the TB outbreak. The DFV decreased before the TB date, reached a minimum during the outbreak, and increased during the recovery phase. As the TB outbreak progressed (during the pre-injury and acute phases), the consumption time (for an equivalent amount of feed eaten) decreased and remained low during the recovery phase. This study confirms that a change in feeding behaviour in a pen can indicate future TB. This also confirms the gradual change of the feeding behaviour over time, reaching its maximum at the TB date.

In addition, Wallenbeck et al.’s statistical analysis of the Swedish data set revealed a significant decrease in DFV 42 to 63 days before the TB event, when compared to matched CTL pens [20]. The DFC was also always reduced in TB pens compared to CTL pens. This difference in feeding behaviour between CTL and TB pens must have been noticed in the current analysis by the ML models, which never predicts TB class in CTL pens. However, both studies did not have the same reference. The Wallenbeck study compared TB pens to matching CTL pens in an attempt to identify eating behaviour predictive of a future TB occurrence [20]. The present analysis used each pen as its own control, as a TB pen was categorised in the TB class when inside the TB window but in CTL when outside. The model could not only detect feeding behaviour that is typical of a future TB event but also changes in feeding behaviour over time that are indicative of TB events. Indeed, a drastic change in feeding behaviour is also indicative of an upcoming TB event [18, 19]. Furthermore, the current approach takes into account a combination of three feeding behaviour metrics (DFV, DFC, and StdFC), improving the likelihood of detecting TB episodes. In a comprehensive book chapter on TB, Valros describes the gradual change in feeding behaviour until the TB date [4]. Daily Feeder Visit can begin to decline months to weeks before the TB date, but DFC appears to be impacted just six days before a TB event [4]. As a result, combining these two indications should increase the model’s accuracy.

## Conclusion and future work

The sensitivity and specificity of certain models (e.g. RF) are very promising, but prediction performances (especially sensitivity) could still be improved using an ensemble model for binary class classification following the model developed by Iwendi et al.[34]. In addition, the same data framework, i.e., a 14 days analysis window combined with a [-35-10] days TB window could be analysed by an elaborate neural network model like long short-term memory recurrent neural network for improved sensitivity. Furthermore, an increased sensitivity rate could potentially be achieved by combining predictions using feeding behaviour data with predictions using other data sources, such as tail position.

In conclusion, a ML model can be deployed in farms with automatic feeders to detect early indicators of TB behaviour at least 35 days before the actual TB event. Thanks to these early warnings, farmers could implement measures to prevent the occurrence of the TB event—for example, by adding more straw as occupational material. Farmers could also start a TB risk analysis to identify the reasons why pigs are disturbed. Continuous implementation of the model on farms would also lead to improved prevention of TB events, serving the welfare of the pigs and bringing an economic benefit to the farmers. Finally, this ML approach could also be a useful tool by allowing the systematic study of the effectiveness of different intervention strategies under controlled conditions.

